# Leveraging collective regulatory effects of long-range DNA methylations to predict gene expressions and estimate their effects on phenotypes in cancer

**DOI:** 10.1101/472589

**Authors:** Soyeon Kim, Hyun Jung Park, Xiangqin Cui, Degui Zhi

## Abstract

DNA methylation of various genomic regions plays an important role in regulating gene expression in diverse biological contexts. However, most genome-wide studies have focused on the effect of 1) methylation in *cis*, not in *trans* and 2) a single CpG, not the collective effects of multiple CpGs, on gene expression. In this study, we developed a statistical machine learning model, geneEXPLORER (<u>gene</u> <u>ex</u>pression <u>p</u>rediction by <u>lo</u>ng-<u>r</u>ange <u>e</u>pigenetic regulation), that quantifies the collective effects of both *cis*- and *trans*- methylations on gene expression. By applying geneEXPLORER to The Cancer Genome Atlas (TCGA) breast and lung cancer data, we found that most genes are affected by methylations of as much as 10Mb from promoter regions or more, and the long-range methylation explains 50% of the variation in gene expression on average, far greater than *cis*-methylation. The highly predictive genes are related to breast cancer, especially oncogenes and suppressor genes. Further, the predicted gene expressions could predict clinical phenotypes such as breast tumor status and estrogen receptor status (AUC=0.999, 0.94 respectively) as accurately as the measured gene expression levels. These results suggest that geneEXPLORER provides a means for accurate imputation of gene expression, which can be further used to predict clinical phenotypes.

## INTRODUCTION

DNA methylation, an essential epigenetic marker, plays an important role in regulating gene expression (1). Methylation within the gene promoter inhibits transcription of the gene (2,3). Methylation in the gene body can be positively correlated with the gene expression level (4). Enhancer regions are associated with low levels of CpG methylation (5). In addition, expression quantitative trait methylations (eQTMs) have found associations between *cis* (regulating transcription of neighboring genes) methylation regions and gene expression (6,7).

In cancer, hypomethylation and hypermethylation were observed at some promoters (8,9). Tumor suppressor genes are inactivated by hypermethylation in promoter regions (9). While aberrant methylation in promoter regions mostly affects transcription in cancer, hypermethylation in gene body regions may not have a noticeable effect on transcription in cancer (10).

Recent studies have examined the effect of methylation in *cis* enhancer regions of genes in cancer. Aran et al.(11) computationally found that methylation in enhancer regions regulates genes more strongly than methylation in promoter regions in cancer, demonstrating the importance of enhancer methylation. Yao et al. (12) inferred cancer-specific *cis* enhancers from methylome and transcriptome analysis in multiple cancer types. However, their studies have focused on the effect of methylation *in cis* (ex. within 1 Mb from Transcription Start Site (TSS) or nearby genes from a CpG site) on gene expression.

To better understand the regulatory role of methylation, studying *trans* (regulating transcription of distant genes) regions is critical. This is because enhancers play an important role in dysregulation of gene expression in cancer (13), and they can be located more than few Mb from a gene (14). For example, a super-enhancer of the MYC gene is reported to be located 1.47Mb from the TSS of the gene in T cell acute lymphoblastic leukemia (15).

In addition, to fully understand the effect of distal regulatory methylation, it is important to consider the collective effect of multiple associated methylations on gene expression, because multiple enhancers regulate expression of a single gene (14,16,17). However, most statistical approaches are limited to testing a single probe and a single gene at a time, such as eQTMs and ELMER (12), making it difficult to quantify the collective effect of CpG methylation on gene expression.

To address these issues, we developed geneEXPLORER (<u>gene</u> <u>ex</u>pression <u>p</u>rediction through <u>lo</u>ng-<u>r</u>ange <u>e</u>pigenetic <u>r</u>egulation), a statistical machine learning method. For each gene, geneEXPLORER identifies CpG methylations, both *cis* and *trans*, that are associated with the gene expression and quantifies the collective effects of multiple CpG methylations. Based on the associated methylation probes, geneEXPLORER builds a predictive model for gene expression. We predicted expression levels of ∼14,000 genes using geneEXPLORER in TCGA breast cancer data and validated the predictions in another breast cancer cohort. We also showed the applicability of geneEXPLORER method to lung cancer. To evaluate the applicability of the gene expressions predicted by geneEXPLORER to downstream tasks, we further predicted the breast cancer phenotypes, such as breast tumor or normal status, estrogen-receptor (ER) status, 5-year survival, and breast cancer subtypes. Since the predicted gene expression represents a methylation portion of the epigenetically regulated gene expression, the present study provides a mechanistic insight into the epigenetic regulation of gene expression and epigenetic effects on cancer phenotypes through gene expression.

## MATERIAL AND METHODS

### DNA methylation and RNA sequencing data from TCGA Breast cancer

To predict gene expression from methylation data, we analyzed TCGA breast cancer data for 873 samples, whose 450K methylation array data and Hi-Seq 2000 gene expression data were available. Among these samples, 788 samples are tumor and 85 samples are normal. The two datasets were downloaded from Xena Public Data Hubs.

### Pre-processing

The values of methylation data in the data hubs are beta values. We transferred beta values to M values because M values are more suitable (closer to normal distribution) for linear regression. Among 485,577 probes, we removed 90,007 methylation probes whose values were missing in more than 20% of the samples. Then, we imputed 31,700 methylation probes whose missing rates were less than 20% using K-means clustering (R package REMP).

For gene expression data, among 20,530 genes, we excluded 3,417 genes whose average expression levels are less than 1 (log2(RPKM+1)) from the prediction. Among the 17,113 genes, TSS sites are available for 16,681 genes from UCSC genome browser. Among these, there was at least one probe in promoter regions for 13,982 genes. We included these genes in our final analysis.

### geneEXPLORER (<u>gene</u> <u>ex</u>pression <u>p</u>rediction by <u>lo</u>ng-<u>r</u>ange <u>e</u>pigenetic <u>r</u>egulation)

In detail, for each gene, we built a linear regression model to predict gene expression using long-range methylation probes.

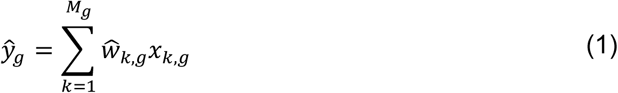

where *ŷ*_*g*_ is the predicted expression of gene *g, x*_*k,g*_ is k-th methylation probe for gene *g, ŵ*_*k,g*_ is the regression coefficient of the methylation probe, *M*_*g*_ is the number of methylation probes within a defined region (e.g. 10Mb or the entire chromosome). To estimate the weight *ŵ* _*k,g*_, we used the elastic-net penalty (18) with α=0.5 (the combination of half Lasso and half ridge penalty) and the penalty was selected through cross-validation using the R package glmnet.

Elastic-net was chosen to predict gene expression using long-range methylations for the following reasons. First, the elastic-net works well with a high-dimensional methylation dataset. Up to ∼38,000 probes were included in the model while the number of samples was only 873. It is impossible to accurately predict gene expression using such a high dimensional data using a model based on regular linear regression models. Second, the elastic-net automatically selects important variables that are associated with a response. By utilizing the elastic-net, geneEXPLORER automatically selects methylation probes that are associated with gene expression from tens of thousands of probes and builds gene expression prediction models based on the probes. Third, the elastic-net works well in highly correlated datasets. Since some of the methylation values are highly correlated due to biological interactions, it is expected that the elastic-net works better than Lasso (18–20).

### Measuring prediction accuracy

To measure prediction accuracy, 10-fold cross-validation (CV) was used. 9 folds of data were used to build a model. The model used methylation values in the remaining fold to predict gene expression. We repeated the procedure 10 times until all gene expressions were predicted. For 81 patients, more than 2 samples existed for the same patient in the dataset (79 patients – 2 samples, 2 patients – 3 samples). We assigned the samples for the same patients to the same fold to avoid a bias. Prediction accuracy (R^2^) was measured as the squared Pearson’s correlation coefficient between predicted gene expression and true gene expression.

### Comparing prediction accuracy of different regions in a gene

We defined a promoter region from 2000bp upstream and 0bp downstream of the transcription start site of a gene (21). Gene regions were obtained using R packages IlluminaHumanMethylation450kanno.ilmn12.hg19, which is annotated by Illumina. The gene regions include promoter region, 5′UTR, first exon, gene body, and 3′UTR.

The long-range regions refer to the regions that maximize prediction accuracy using geneEXPLORER. The range is from 1Mb from the promoter region to the entire chromosome on which the gene is located. We fitted the elastic-net model (Eq.1) for each region and each gene. We showed prediction accuracy of 13,910 overlapping genes (among 13,982 genes) for which all the following conditions were satisfied; (a) Gene location was available in the R package (b) there was at least one probe in the promoter region.

### Investigation of various distances from promoter regions of genes

For each gene, we built elastic-net models using methylation CpG sites for various distances (1, 2, …, 10, 20, 30, 40, 50 Mb) from the promoter region of the gene were built. The elastic-net model was also built using all CpG sites on the same chromosome where the gene is located. Prediction accuracy was evaluated using 10-fold CV R^2^. Then, distances were selected that maximized the prediction accuracy for each gene.

### Evaluating prediction accuracy using multiple regression based on *trans* eQTMs

Since traditional multiple regression cannot handle high dimensional data (the number of samples < the number of probes), methylation probes were pre-screened before fitting multiple regression models. For each gene, association between a gene and each methylation probe was tested using single linear regression models, where the covariate is a probe and the response is a gene (expression). Probes were tested in the entire chromosome on which the gene was located. Multiple-testing adjustment was performed for each gene using Bonferroni correction at significance level 0.05. Using the significantly associated probes, we built a multiple linear regression model for each gene. If the significantly associated genes were still more than the number of samples in a training set, a ridge regression model (22) was fitted. 10-fold CV was used to calculate prediction accuracy (CV R^2^).

### Testing on an independent cohort

geneEXPLORER was trained using TCGA breast cancer data and tested on GSE39004 data. geneEXPLORER models were built using TCGA data for each gene, and models were selected that minimized CV error using 10-fold CV. Using the methylation probes from the test dataset as inputs of the models, gene expression was predicted for 13,027 genes. Test R^2^, which is squared Pearson’s correlation coefficient between the predicted gene expression and the observed gene expression of the test dataset, was calculated.

For comparison, multiple regression models based on eQTMs were used, as in the previous section. We used the training data to select significant probes, using univariate tests with Bonferroni correction (*α* = 0.05) and to fit multiple regression models. Using the methylation array data in the test data as input to the models, gene expression was predicted using the multiple regression model for each gene, and test prediction accuracy was calculated. We limited long-range distance to 10Mb from promoter regions to save computational time.

### Applying geneEXPLORER to lung cancer dataset

For lung cancer data analysis, methylation probes and genes were pre-screened in the same way as for breast cancer (395,616 probes and 14,256 genes). The TCGA lung cancer data consist of 856 samples (827 tumor and 29 normal), for which both 450K-based methylation data and RNA-Seq gene expression data were available.

To compare prediction accuracies of gene expression of both types of cancer, test prediction accuracy within each dataset was measured. For each type of cancer, the dataset was divided into a training set (4/5 of the samples) and a test set (1/5 of the samples). The procedure was repeated 5 times until all gene expression data was predicted. Test R^2^ was calculated using the squared correlation coefficient between the predicted gene expression and observed gene expression. The model was trained using methylation probes within 10Mb from the promoter regions of the genes.

### Predicting clinical phenotypes

Breast cancer status and estrogen receptor (ER) status were predicted using the predicted gene expression. For cancer status, 788 samples were tumor cells and 85 samples were normal cells, among 873 samples from the TCGA breast cancer data. For ER status, 632 samples had ER-positive status,183 samples had ER-negative status, while 58 samples had missing ER status.

To predict the clinical phenotypes, 13,982 gene expressions were first predicted in test datasets in the same cohort. The data was divided into a training set (4/5 of the samples) and a test set (1/5 of the samples). Using the training dataset, 10 folds cross-validation (4/50 of samples are in each fold) was used to select a model that maximized prediction accuracy using probes within ±10Mb from the promoter regions. By inputting methylation in the test dataset into the selected model, gene expression in the test dataset was predicted. The procedure was repeated five times until all gene expression data was predicted.

Next, a penalized logistic regression model (elastic-net) was fitted using the 13,982 gene expressions as covariates, and a phenotype as a binary response, as described in the following equation:

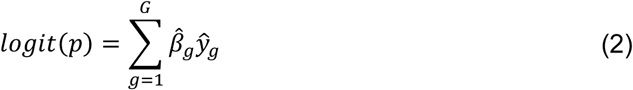

where *p* is the probability of a phenotype to be “Yes” (e.g. tumor/ER-positive), *ŷ*_*g*_ is the predicted expression of gene *g*, 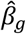 is the regression coefficient of gene *g,* and *G* is the number of predicted genes (13,982).

Note that the elastic-net model automatically selects gene expression that is associated with the phenotype. Prediction accuracy was evaluated by area under the ROC curve (AUC) using 10-folds CV.

## RESULTS

### Gene expression prediction by long-range epigenetic regulation (geneEXPLORER)

geneEXPLORER quantifies the regulatory effects of CpG methylation on gene expression by exploiting long-range regulatory elements up to the entire chromosome on which the gene is located. Because multiple distal regulatory elements interact to regulate gene expression (14,16,17), geneEXPLORER is expected to make more accurate predictions of gene expression than the models that only use *cis-*elements. As gene expression is often profiled to determine clinical phenotypes, the predicted gene expression, therefore, can also be used to predict the phenotypes. The prediction accuracy of phenotypes can also indicate the collective effects of distal methylations on the phenotypes through gene expression regulation.

The training procedure of geneEXPLORER is shown in Figure 1. First, given a training set of methylation data across samples, an elastic-net model (18) was build, geneEXPLORER, where covariates are long-range methylation probes within a certain distance from the promoter region (*L*_*g*_ in Figure 1B) and a response is the observed expression level of a gene (Figure 1C). Elastic-net was chosen because the elastic-net works well in high-dimensional methylation dataset and automatically selects methylation probes that are associated with gene expression. During the training phase, geneEXPLORER identifies methylation CpG sites that are associated with gene expression and estimate the weights of the identified CpG sites. Second, geneEXPLORER with trained weights is used to predict the gene expression using methylation in the test dataset. Then, we measure the prediction accuracy using R^2^. We repeat the procedure for all genes. Next, using the predicted gene expression by geneEXPLORER as an input, we further build elastic-net logistic regression models to predict binary clinical phenotypes (Figure 1D). Since we use predicted genes (p=∼14,000) as covariates, instead of methylation probes (p=∼500,000), it is possible to build the prediction model without suffering due to the very large number of methylation probes. Through the prediction model, we could estimate the effect of methylation on the phenotypes through gene expression regulation.

**Figure 1.**
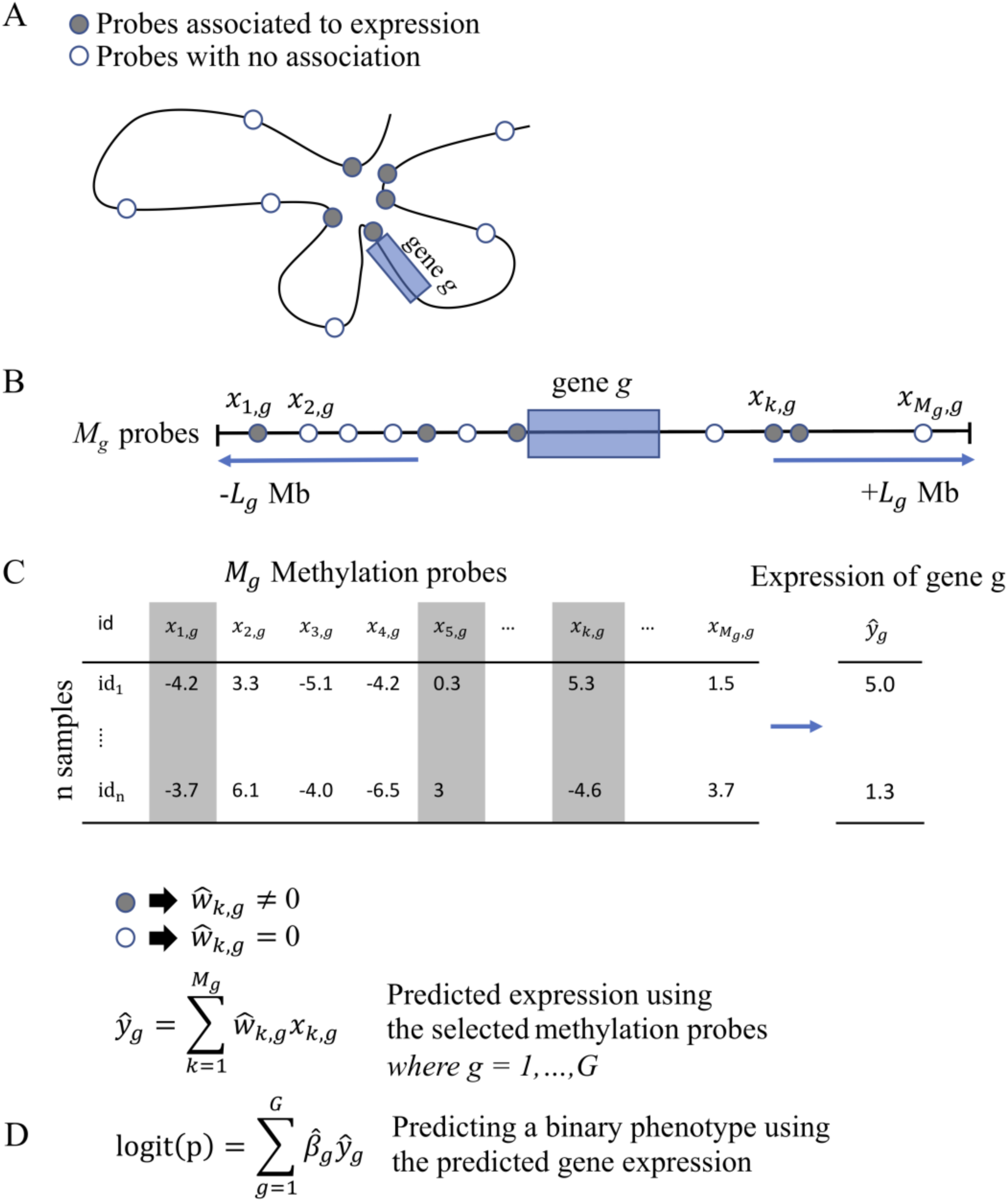
GeneEXPLORER modeling: (A) Several methylation probes are associated with gene expression, and they can be located far from the gene due to chromatin looping structure. (B) Straightened genome upstream and downstream *L*_*g*_ Mb from the promoter region of the gene g. There are *M*_*g*_ numbers of probes in the range. (C) Predicting gene expression from the methylation probes. Methylation data to predict the expression of gene, g consist of n samples and M_g_ probes. The shaded columns are an example of probes that are associated with gene expression. Our model, geneEXPLORER, identifies the associated probes and estimates the weights of them. Gene expression of g is predicted by summing the weighted methylation values. The procedure is repeated for each gene. (D) Application of geneEXPLORER: Predicting phenotypes from the predicted gene expression. After predicting gene expression on the entire genome, we estimated the effects of the predicted regulated gene expression on several binary phenotypes (see Methods).

### The collective effect of long-range methylation on gene expression is higher than that of promoter and gene region methylation on gene expression

First, using 13,910 expressed genes in 873 TCGA breast cancer samples, we investigated how distance of methylation affects gene expression: from ±1Mb from the promoter region to the entire chromosome on which the gene is located (see Methods). As the associated methylation probes were different for each gene, we selected the distance that maximized prediction accuracy (CV R^2^) (Figure 2A, **Figure S1, Figure S2**). For most of the genes, long-range methylation probes were required to predict gene expression accurately: 84% of the genes need methylation probes more than ±10Mb away to achieve the best prediction accuracy. 49% of the genes required including methylation probes more than ± 35 Mb away from the genes to maximize prediction accuracy (Figure 2A). Also, 31% of the genes required methylation values from the entire chromosome to maximize their gene expression accuracy. This shows that most genes can be affected by distal regulatory elements that are located more than 10Mb from the promoter regions. A possible reason is that even though most enhancers are within a few Mb from the regulated gene (14) (also supported by **Figure S1**), there can be still several enhancers that are far away (more than 10 Mb).

**Figure 2.**
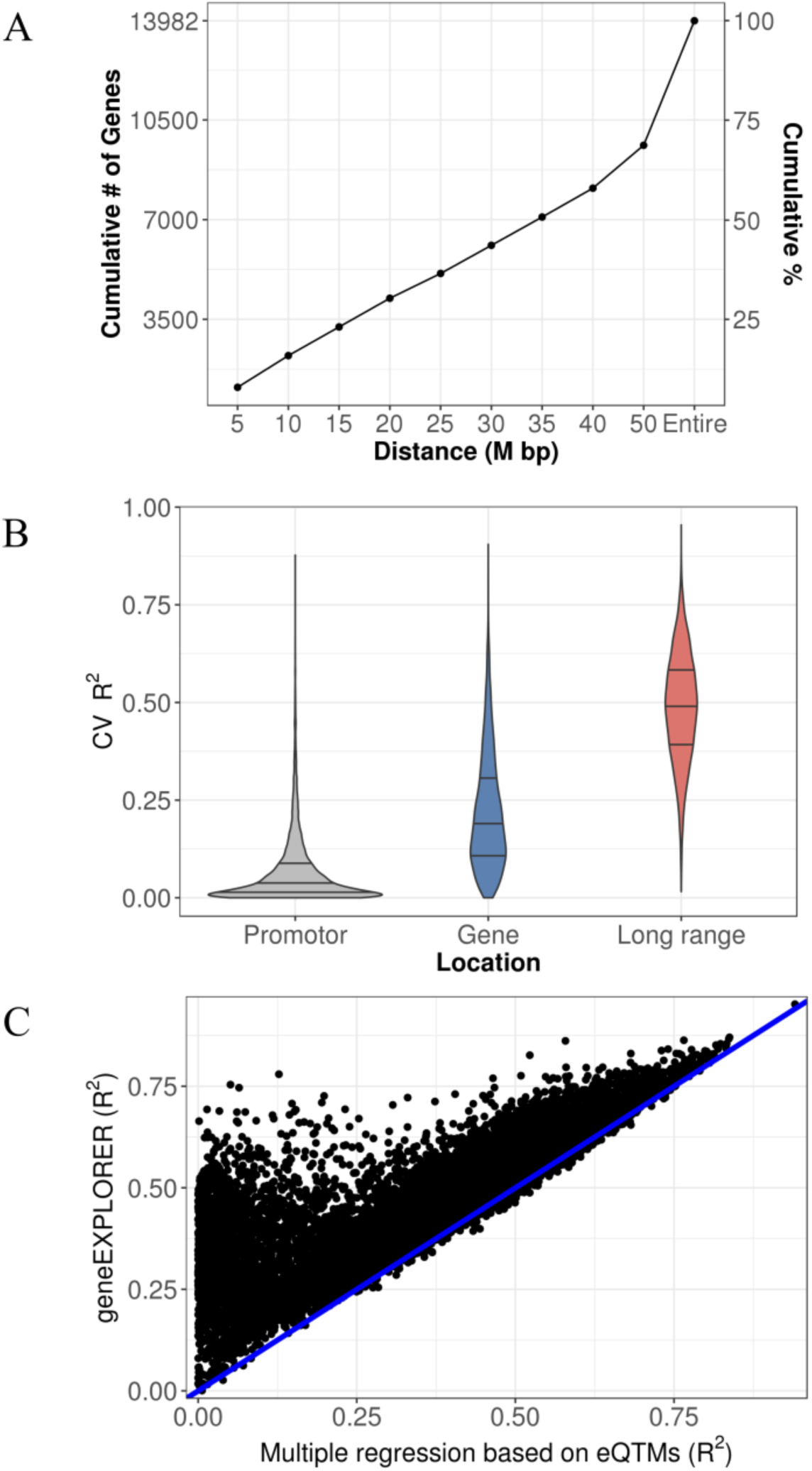
Prediction power comparisons using cross-validation (CV) (A) Distance (from the promoter region) of probes (L_g_ in **The *collective effect of* B**) that maximized prediction accuracy for each gene and the cumulative frequency of the distances and the percentage. The distance was selected from ±1Mb from promoter regions to the entire chromosome on which the gene is located. (B) Gene expression prediction power (CV-R^2^) by region using TCGA breast cancer data: the predictive models were developed based on methylation probes in 1) the promoter, 2) the gene, and 3) long-range regions. We plotted 13,910 genes for which at least one probe is included in the promoter region of the gene. The three lines in the violin plots indicate 25%, 50%, and 75% percent quantiles, respectively. (C) Prediction power (CV R2) comparison: geneEXPLORER vs. multiple regression based on expression quantitative trait methylations (eQTMs) using the entire chromosome. Data points are 13,982 genes. The blue line is y=x.

To understand the methylation effect on regulatory regions, gene expression levels were predicted using methylation probes in 3 different regulatory regions: 1) promoter, 2) gene, 3) long-range regions. Gene regions include the promoter region, 5′UTR, first exon, gene body, and 3′UTR as Illumina annotated.

Methylation in long-range predicts gene expression far better (average CV R^2^=0.486) than methylation in either promoter (average CV R^2^=0.064) or gene regions (average CV R^2^=0.218) (Figure 2B). A possible reason is that the collective effects of *trans*-methylation can exert a stronger effect on gene expression than *cis-*methylation in the promoter or gene region, although individual effects of *trans*-methylation may be weaker than that of *cis*-methylation. These results suggest that distal methylation outside of the promoter and of the gene regions can play more important roles in regulating gene expression than methylation on the promoter and the gene regions.

### Prediction comparison between geneEXPLORER and multiple regression using expression quantitative trait methylations (eQTMs) in TCGA breast cancer

To understand the prediction performance of geneEXPLORER in comparison to a traditional statistical method, the prediction accuracy of geneEXPLORER in predicting gene expressions was compared to a multiple regression model based on trans-eQTMs. For eQTMs, probes are selected by univariate tests with Bonferroni correction (p-value < 0.05) for each gene using methylation probes in the entire chromosome on which each gene is located (see Method). With methylation probes in the same range, geneEXPLORER outperformed the multiple regression model based on eQTMs (Figure 2C). geneEXPLORER predicted 97% of the gene expressions (13,569 out of 13,982) better than eQTMs. A possible reason may be that multiple testing correction methods in eQTMs tend to be too conservative to control false positives, thus weakening the power to detect significant probes that are associated with a gene. Too few true positive probes in the multiple regression models make impossible to predict gene expressions better than geneEXPLORER, which automatically selects probes without statistical tests.

### Testing geneEXPLORER on an independent cohort

To show that geneEXPLORER can be used to predict gene expression in an independent cohort, geneEXPLORER trained in the TCGA BRCA was tested on an independent breast cancer cohort. This dataset consists of methylation 450K array and gene expression microarray datasets of 57 breast tumor samples and 8 adjacent normal samples (GSE39004). The result was compared with that of the multiple regressions based on eQTMs for 13,027 expressed genes. We found that, for a majority of the genes (10,189, 78%), geneEXPLORER predicted gene expression better than the multiple regression based on eQTMs in the independent cohort (Figure 3), demonstrating its applicability to independent datasets of the same cancer type.

**Figure 3.**
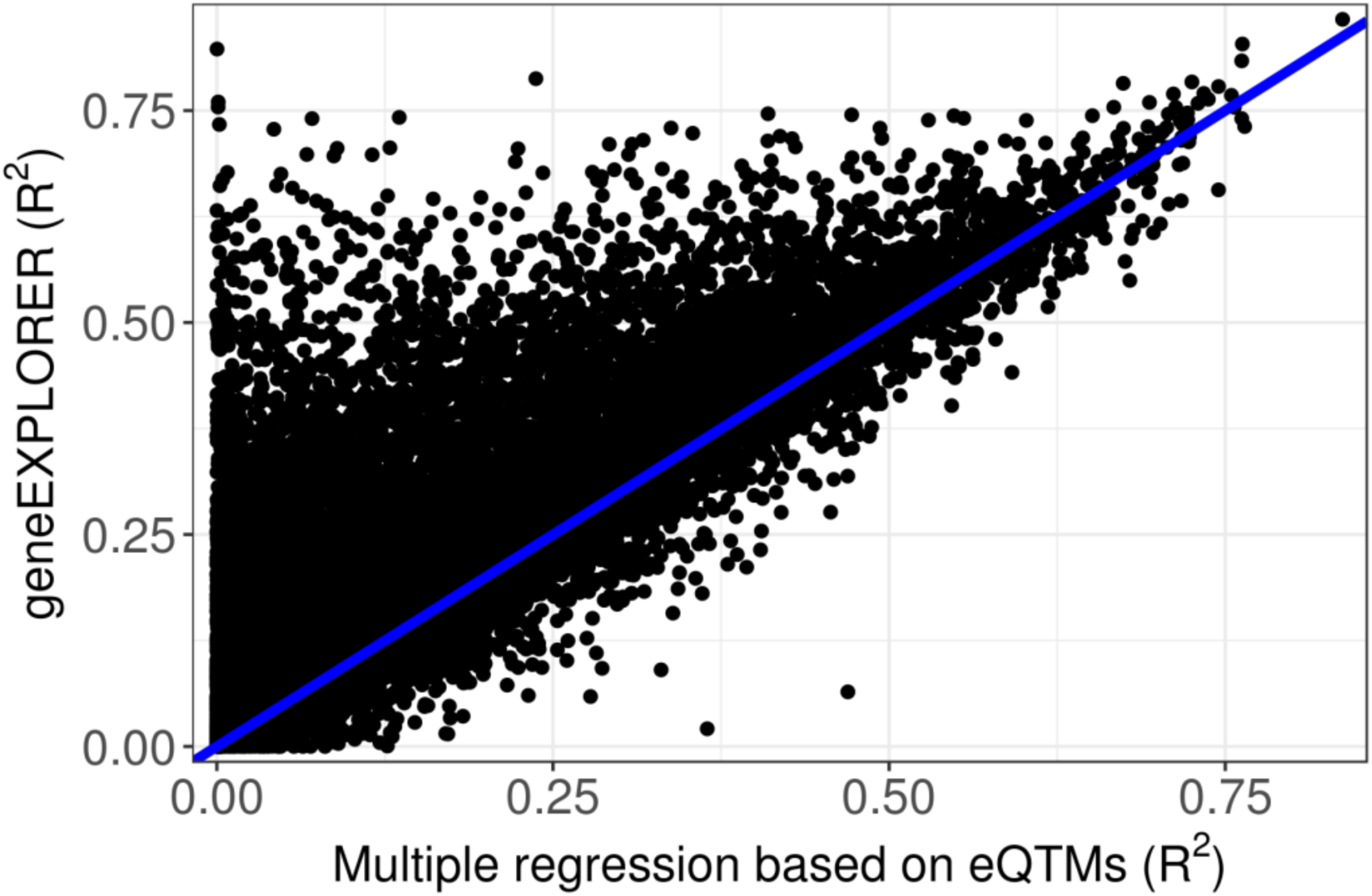
Prediction accuracy on an independent breast cancer cohort (GSE39004). Using the prediction model trained on TCGA breast cancer, prediction accuracy tested on GSE39004 data was compared between geneEXPLORER and a multiple regression based on eQTMs. Methylation probes in ±10Mb from the promoter regions were used for 13,027 genes.

### Applicability of geneEXPLORER to another type of cancer

To demonstrate its applicability to other types of human cancer, geneEXPLORER was applied to l ung cancer. TCGA lung cancer data is the combination of lung adenocarcinoma and lung squamous cell carcinoma (n=856 samples). We trained and tested the model for each cancer type (see Methods). We compared R^2^ for breast cancer data and lung cancer data for 11,665 overlapping genes (Figure 4). Lung cancer showed a similar high prediction accuracy as the breast cancer (R^2^ 0.441 for breast cancer and 0.428 for lung cancer). This demonstrated that geneEXPLORER can b e applied to other cancer types to predict gene expression in the presence of methylation data.

**Figure 4.**
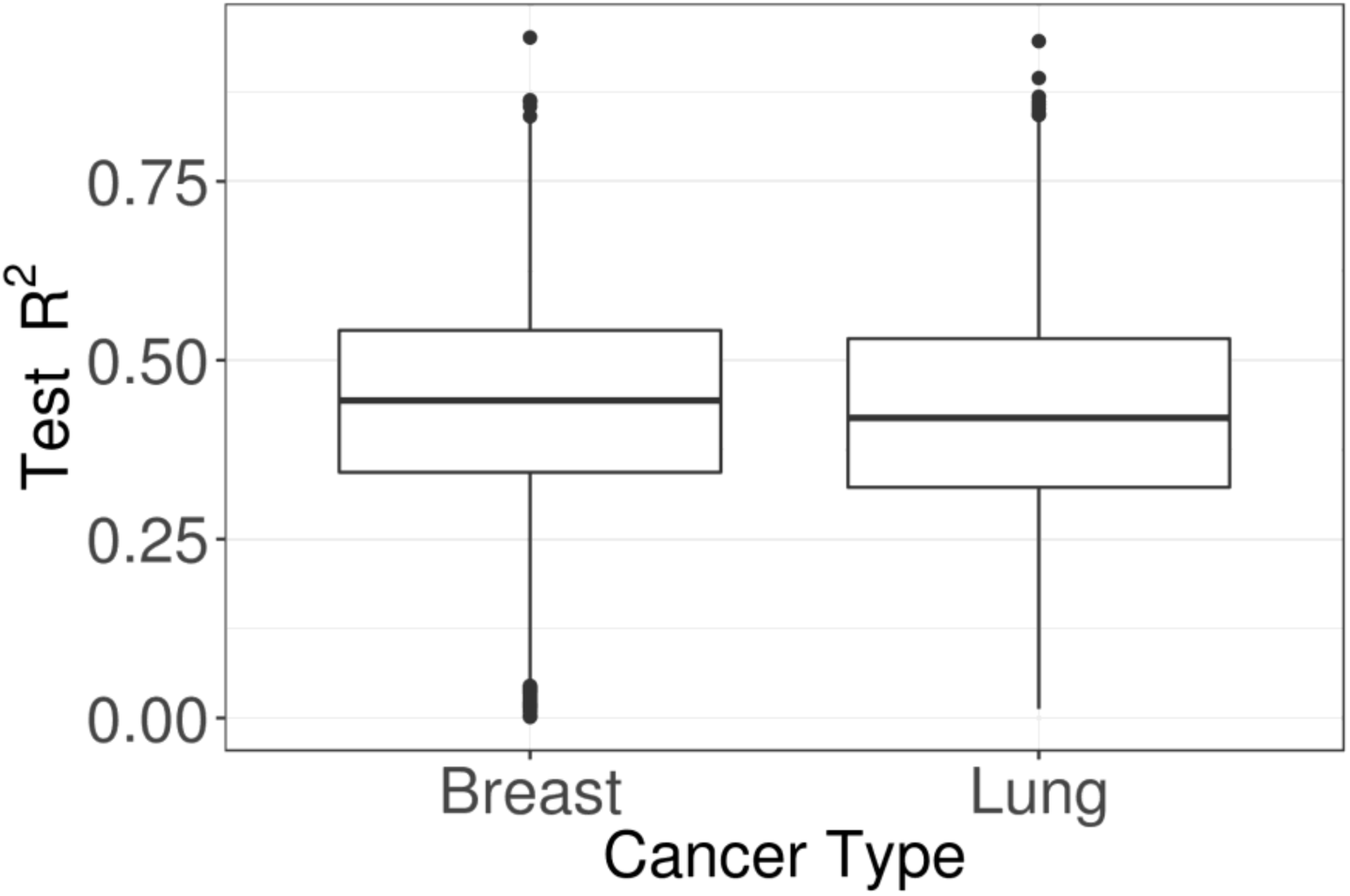
Boxplot of test R^2^ for TCGA breast and lung cancer data. geneEXPLORER for TCGA lung cancer data demonstrated a similarly good prediction accuracy as for TCGA breast cancer. The result was shown for overlapping 11,665 genes.

### geneEXPLORER accurately predicts expression of tumor-associated genes

We found that geneEXPLORER accurately predicts expression of multiple genes which play important roles in breast cancer. Examples are shown in Figure 5. Polymorphisms of GSTT1, the highest predicted gene, are established risk factors for breast cancer (23–25). The mutation of GATA3 is known to lead to luminal tumors (26). ESR1 is the estrogen-receptor gene, common in primary breast cancers, whose mutation is indicative of resistance to anti-estrogen therapies (27–32). In addition, breast cancer risk–associated SNPs are enriched in the cistromes of FOXA1 and ESR1 (33). High expression of SOX10 is observed in triple-negative and metaplastic breast carcinomas (34). ERBB2 is a well-known oncogene of breast cancer (35).

**Figure 5.**
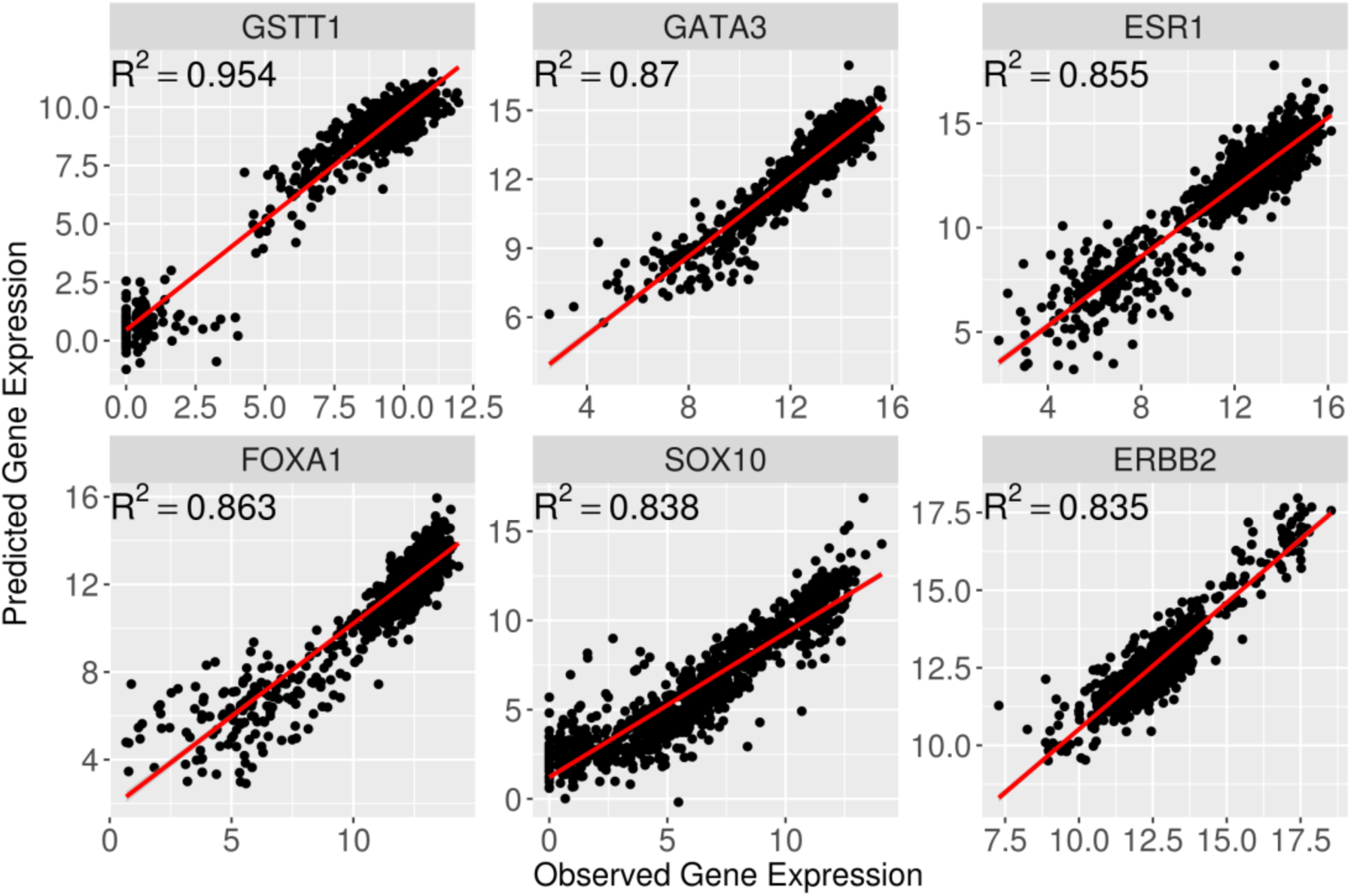
Examples of highly predicted genes that are associated with Breast cancer. The genes were predicted by geneEXPLORER using TCGA breast cancer data. R^2^ is Cross-validation prediction accuracy.

In addition, we also found that geneEXPLORER predicted many oncogenes and tumor suppressor genes with high prediction accuracy (Table 1). This means that those genes are also regulated by long-range methylation. Since many abnormal enhancer activities are found in cancer and enhancer regions are often hypomethylated (13), the oncogenic mechanism involving the oncogenes and tumor suppressor genes can be associated with abnormal activities in methylation. The roles of these genes in breast cancer have been widely studied at the genetic or transcriptomic level but not as much in epigenetics. Since methylation through long-range interactions predicted a substantial part of gene expression, geneEXPLORER can further help to discover the tumorigenic role of long-range methylation in human cancer.

**Table 1.**
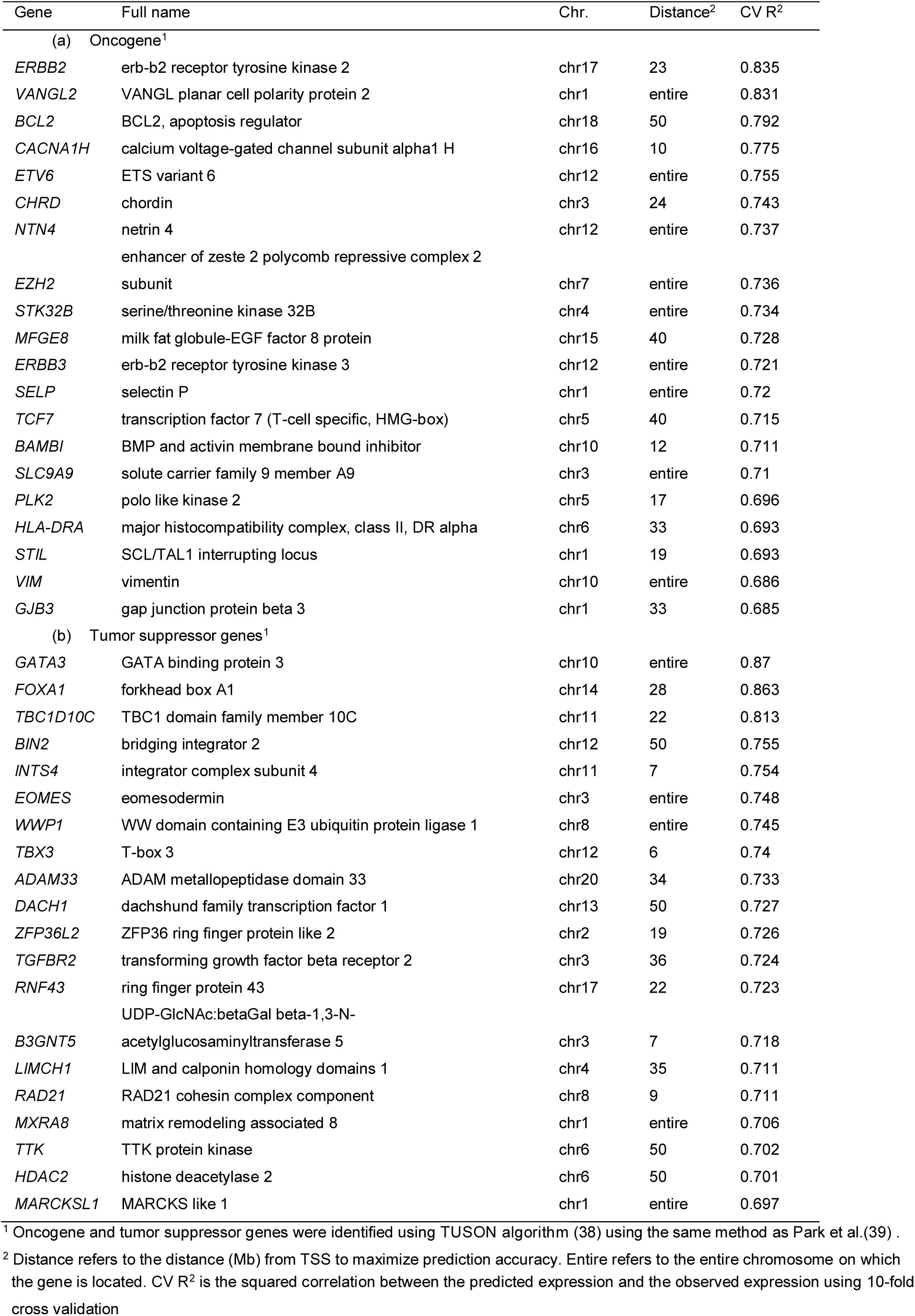
Best predicted 20 oncogenes and 20 tumor suppressor genes by geneEXPLORER

### geneEXPLORER accurately predicts clinical features of human cancer based on the predicted gene expressions

Since gene expression profiles often reflect clinical phenotypes (36), to determine potential clinical applications of geneEXPLORER, we built predictive models using the predicted gene expressions to predict clinical phenotypes of TCGA breast cancer data (see Methods). Based on the predicted expression levels of 13,982 genes, we predicted cancer status (tumor / normal), Estrogen Receptor (ER) status (positive / negative), 5-year survival (yes / no) and PAM50 breast cancer subtypes. Due to the high prediction accuracy of the breast cancer-related genes, high prediction accuracies of these phenotypes were expected.

Consistent with the expectation, by comparing prediction accuracy between the model using the predicted gene expressions and the model using the observed gene expression, we found that virtually no difference between the predicted gene expressions and the observed gene expressions in predicting the phenotypes (Figure 6, **Figure S5**, and **Table S1**). Notably, gene expression predicted by methylation almost perfectly predicted both cancer status and ER status (AUC=0.999 and 0.94 respectively) (Figure 6).

**Figure 6.**
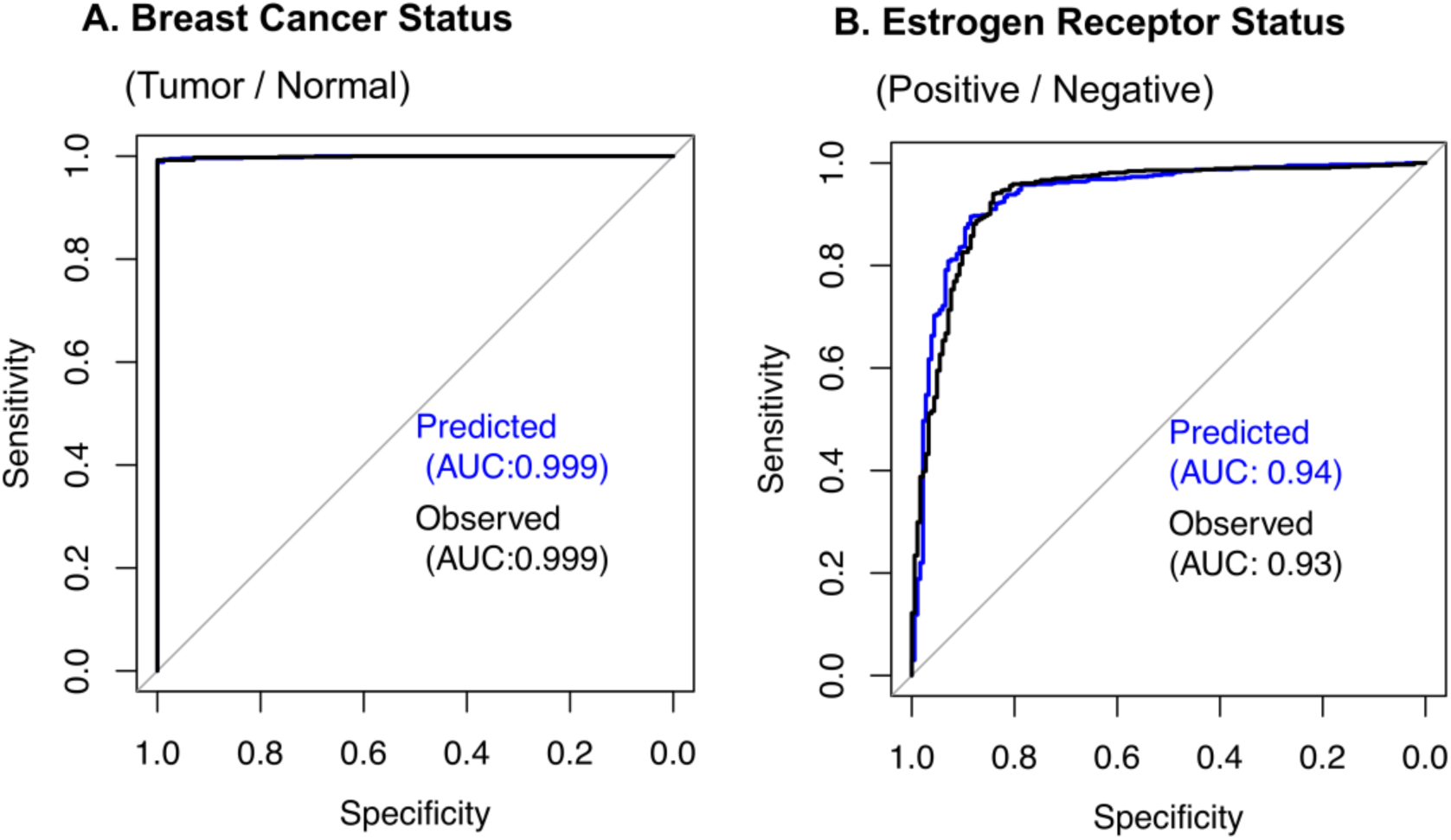
ROC curve for predicting clinical phenotypes using the gene expression predicted by geneEXPLORER (predicted) vs. observed gene expressions (observed): the predicted gene expression predicts the phenotypes as good as the observed gene expressions with perfect prediction accuracy.

Since the predicted gene expression was the portion of gene expression regulated by methylation, the high prediction accuracy of the clinical features implies that long-range methylation plays a critical role in determining the phenotypes through regulating gene expressions in breast cancer. This shows that the predicted gene expression can be applied to help diagnose cancer phenotypes or develop personalized treatments as was the approach using observed gene expressions (20), even when gene expression data are not available.

## DISCUSSION

In this paper, we developed a statistical machine learning model, geneEXPLORER, to quantify methylation effects on the gene expression. Methylation of both *cis-* and *trans-* CpG sites was incorporated into the statistical model and the methylation effect of not only a single CpG site but also the collective effects of long-range CpG sites was measured. Applying geneEXPLORER to the TCGA breast cancer dataset demonstrated that 1) most genes are affected by methylation more than 10Mb from promoter regions; 2) long-range methylation highly affect gene expression, far greater than the effect of methylation in the promoter regions or gene body regions; 3) geneEXPLORER outperformed multiple regression models based on eQTMs for the most highly expressed genes in TCGA breast cancer datasets as well as an independent cohort; 4) many highly predicted genes were related to breast cancer, such as oncogenes and tumor suppressor genes; 5) the predicted gene expression predicted breast cancer status and estrogen receptor status with almost perfect prediction accuracy, where the predicted gene expression and the observed gene expression predicted the phenotypes equally well.

geneEXPLORER was partly motivated by Gamazon et al. (19) who predicted gene expression using SNPs nearby to the genes. However, their models showed a markedly lower prediction accuracy than geneEXPLORER (mean CV R^2^=0.15 vs mean CV R^2^=0.486). The lower accuracy could be due to smaller effects of SNPs as opposed to effects of methylation on gene expression, due to smaller genomic regions considered (1Mb from TSS), or different tissue and disease types. Also, Gamazon et al did not directly use the predicted gene expression levels to predict phenotypes. Rather, they developed a method called prediXcan to test the association between the predicted gene expression and several phenotypes. In this study, we used the predicted gene expression to predict clinical phenotypes, showing strong effects of methylation on phenotypes through gene expression regulation.

geneEXPLORER showed much better gene expression prediction accuracy compared to our previous model, MethylXcan (37), which only incorporated CpG sites in the gene region (from promoter to 3’UTR regions) to predict gene expression. In the MethylXcan study, average CV R^2^ were 0.05 and 0.08 in two datasets while geneEXPLORER showed average CV R^2^ of 0.49 in TCGA breast cancer. While the difference can be partly attributed to different penalization methods (Lasso for MethylXcan vs. elastic net for geneEXPLORER), different tissue/diseases (PBMC and adipose in normal or atopic asthma patients for MethylXcan vs. breast cancer for geneEXPLORER), these results suggested that the major difference arises from incorporating long-range methylation in geneEXPLORER while MethylXcan only used gene regions, which is consistent with the result in Figure 2.

geneEXPLORER could not be tested on an independent dataset with the same platform on which it was trained – geneEXPLORER was trained using RNA-seq data but it was tested using gene expression array data (Figure 3). The reason is publicly available datasets with 450K methylation array and RNA sequencing in breast cancer were not available with sufficient sample size. Since only a dataset with 450K methylation array and gene expression array for breast cancer patients (GSE39004) was found, geneEXPLORER on this dataset was tested. This showed worse prediction accuracy than when it was tested within the RNA-seq data (RNA-seq: R^2^=0.444 vs microarray: R^2^=0.263; **Figure S3**), maybe due to the difference between array data and sequencing data, in addition to fitting bias between the training set and the test set.

We showed the applicability of geneEXPLORER in another cancer type, lung cancer (Figure 4). The model was trained in lung cancer and tested in the same cancer type. The prediction accuracy of gene expression was as high as that in breast cancer. This implies that geneEXPLORER method can be applied to any kind of cancer. However, one caution is that the model should be trained in a cancer-specific manner as we showed in **Figure S4** since enhancers are cancer-specific (13).

The scope of this study was limited to predicting gene expression and not identifying/discovering regulatory elements such as enhancers. However, since geneEXPLORER selects CpG sites that are associated with gene expressions, the selected CpG sites could be in enhancer or insulator regions. Therefore, geneEXPLORER may be further developed to identify regulatory regions with stability selection approaches (20).

In conclusion, we developed geneEXPLORER, which identified methylation probes that regulate gene expression using *cis-* and *trans*-methylation. To the best of our knowledge, geneEXPLORER is one of the first to estimate the collective *cis-* and *trans*-effects of methylation on gene expression. Using geneEXPLORER, we found that the collective *trans*-effects are greater than *cis-*effects of methylation. geneEXPLORER predicted about half of gene expression variations on average, which was far more accurate than the estimation using genetic variants from Gamazon et al. (19). In addition, the predicted epigenetically regulated gene expression successfully predicted cancer phenotypes such as cancer and ER receptor status as accurate as the observed gene expressions. Given these results, future application of geneEXPLORER can be 1) imputation of gene expression for other cancer types or other diseases, 2) discovery of regulatory elements, and 3) diagnosis of disease and prediction of phenotypes.

## Supporting information

Supplementary Data are available at NAR online.

## ACKNOWLEDGEMENT

We thank Mr. Irmi Willcockson for English edits and Xin Gene for helping us to go through IRB process.

## FUNDING

This work was supported by US National Institute of Health: [T32 HL 129949 to S.K.]; and Cancer Prevention Research Institute of Texas [RP170668 to D.Z.]. Funding for open access charge:

### Computational Resources

USCS genome browser https://genome.ucsc.edu/

TCGA breast cancer data from UCSC XENA

https://xenabrowser.net/datapages/?cohort=TCGA%20Breast%20Cancer%20(BRCA)&removeHub=https%3A%2F%2Fxena.treehouse.gi.ucsc.edu%3A443

TCGA lung cancer data from UCSC XENA

https://xenabrowser.net/datapages/?cohort=TCGA%20Lung%20Cancer%20(LUNG)&removeHub=https%3A%2F%2Fxena.treehouse.gi.ucsc.edu%3A443

Gene expression omnibus GSE39004 dataset

https://www.ncbi.nlm.nih.gov/geo/query/acc.cgi?acc=GSE39004

## AVAILABILITY

geneEXPLORER is an open source collaborative initiative available in the GitHub repository (https://github.com/SoyeonKimStat/geneEXPLORER)

