## Supplementary material for "Leveraging collective regulatory effects of long-range DNA methylations to predict gene expressions and estimate their effects on phenotypes in cancer"

**SUPPLEMENTARY DATA**

**Long-range methylation affects expression of genes**

We investigated how distantly methylation affects gene expression from 2Mb from the promoter region to entire chromosome on which the gene is located (**Figure S1**). While methylation probes within 2Mb account for 91% of prediction accuracy compared to methylation probes in the entire chromosome, multiple probes affect gene expression from more than 50Mb from the promoter region. This implies that some enhancers can be located very far from genes, in contrast to current thought that enhancers are always located within few Kb to few Mb from a gene (13,14).

The distance that maximized prediction accuracy for each gene was also selected. This (average $R^2$=0.486) improved geneEXPLORER by 14% compared to the model that used methylation probes within 2Mb. CV selected different distances for different genes, implying that enhancers were located at various distances for different genes (**Figure S2**). However, for most of the genes, methylation sites far from the genes were required to predict gene expression accurately: 84% of genes required methylation probes more than 10Mb distant for the best prediction and 31% of genes needed methylation probes in the entire genome to achieve best prediction accuracy. This implies multiple enhancers are far away from the genes for most genes.

**Comparison of prediction accuracy using geneEXPLORER between NGS and microarray platforms**

We compared the performance of gene expression prediction between next generation sequencing (NGS) and microarray, even though geneEXPLORER was trained using gene expression using NGS (**Figure S3**). For NGS prediction, since a breast cancer dataset for which both NGS and 450K methylation array was not available, TCGA breast cancer data was divided into training (4/5 of the samples) and test (1/5 of the samples) datasets. A model was trained and selected using CV (using methylation probes within 10Mb from promoter regions) within the training set and prediction accuracy of test dataset was measured. The procedure was repeated 5 times until all samples were imputed. For microarray prediction, geneEXPLORER was trained using the TCGA breast cancer dataset and tested on another breast cancer dataset (GSE39004), which had both 450K methylation array and microarray gene expression.

As expected, geneEXPLORER predicted NGS gene expression much better than microarray gene expression**:** on average, test $R^2$ of NGS was 0.444 while test $R^2$ of microarray is 0.263. However, it was also able to predict gene expression with moderate prediction accuracy (the average correlation coefficient was 0.514).

**Cancer specificity of geneEXPLORER**

Since enhancers are cancer specific, we expected that geneEXPLORER models are cancer specific as well. To demonstrate this, we applied geneEXPLORER trained in TCGA breast cancer data to predict 13,823 genes of TCGA lung cancer data. The results demonstrated that the model did not work in lung cancer (mean $R^2$=0.02), confirming our hypothesis (**Figure S4**).

**Predicting additional phenotypes using predicted gene expression**

In addition to cancer status and ER status, 5 years survival and breast cancer sub-type were also predicted. For survival data, since 732 samples (83.8%) were censored among 873 patients, there were only 298 samples whose 5 year-survival data are available. 207 patients died before 5 years and 91 patients lived more than 5 years. If censoring occurred before 5 year of follow-up, 5-year survival is indicated as NA. If censoring occurred after 5 year of follow-up, 5-year survival is indicated as Yes. The model predicted 5 years survival with lower accuracy (AUC=0.71) than cancer status or ER status, possibly due to the high portion of censored data (**Figure S5**).

The breast cancer sub-type status was available for 620 samples. The subtypes are Luminal A (LumA), Luminal B (LumB), Triple-negative/basal-like (Basal), HER2-enriched (Her2), and Normal-like (Normal). Using the predicted gene expression, breast cancer subtypes were accurately predicted with 0.174 mis-classification error. Using observed (true) gene expression, breast cancer subtype was predicted with similar prediction accuracy with 0.134 mis-classification error (**Table S1**).

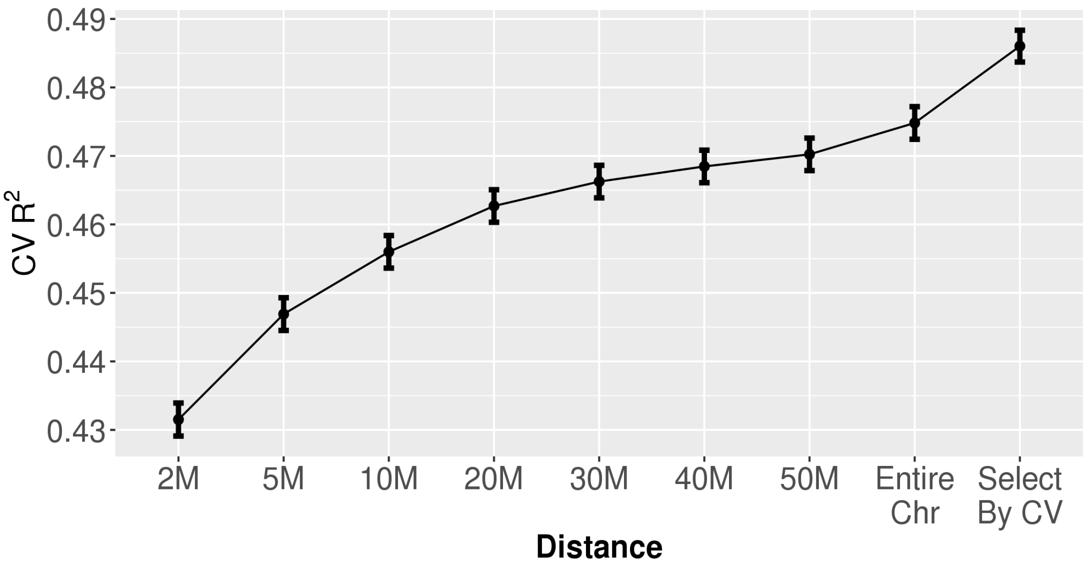


**Figure S1. Improving prediction accuracy by extending distance from promoter regions**: More probes candidate enhancer probes were considered by including all probes within 2Mb, 5Mb,10Mb, 20Mb,30Mb,40Mb, and 50Mb from promoter regions as inputs for the model. Finally, all probes in the entire chromosome on which the gene located were included. Further, we selected the distance which maximized prediction accuracy by CV for each gene and included all probes within the distance as inputs for the model. 13,982 genes were predicted. The dots are means and bars are 2 standard errors.

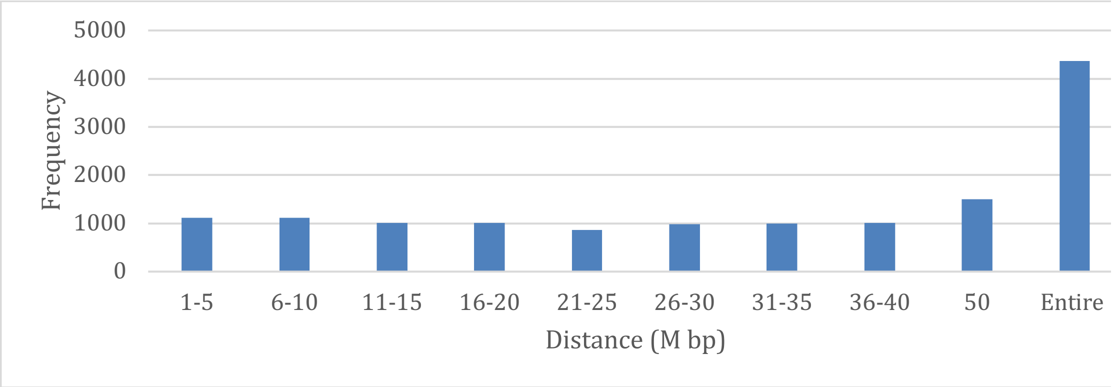


**Figure S2. The distances from promoters which predict gene expression the best: frequency is the number of genes.** To maximize prediction accuracy, most genes require inclusion of methylation very distant to the genes.

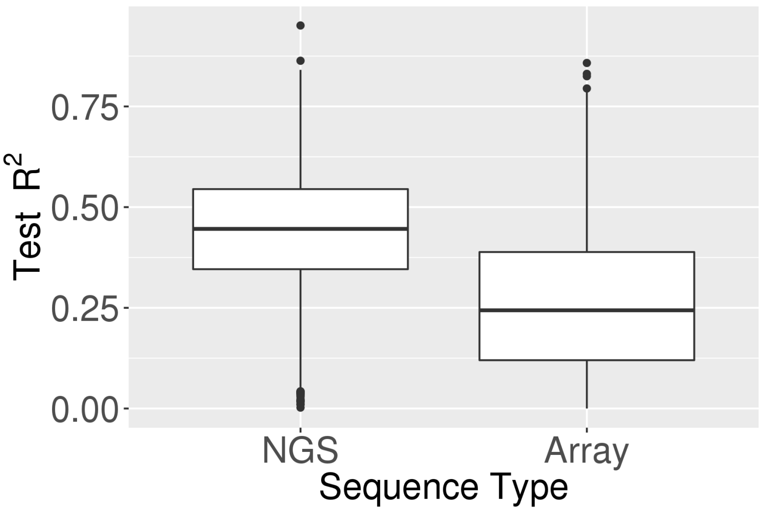


**Figure S3**. **Comparison of prediction accuracy using geneEXPLORER between NGS and microarray platforms**: geneEXPLORER trained on TCGA breast cancer and tested prediction accuracy of gene expression on gene expression data using NGS methods (Left) and using microarray method (Right) The results demonstrate prediction accuracy of 10,972 overlapping genes in both datasets.

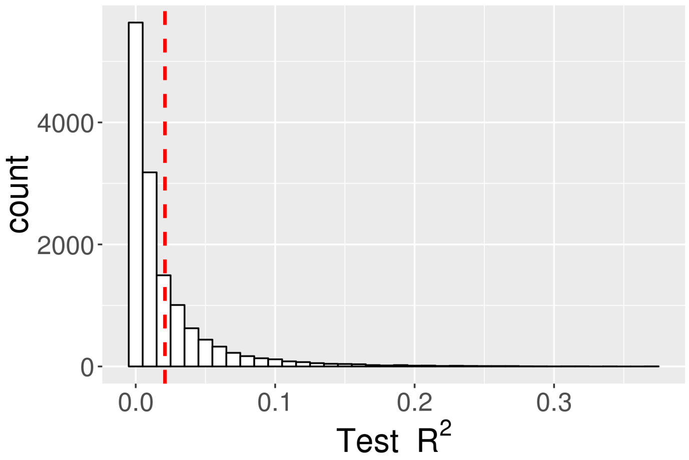

**Figure S4. Cancer specificity of geneEXPLORER**. When gene EXPLORER trained on breast cancer was tested in lung cancer, it showed low prediction accuracy.

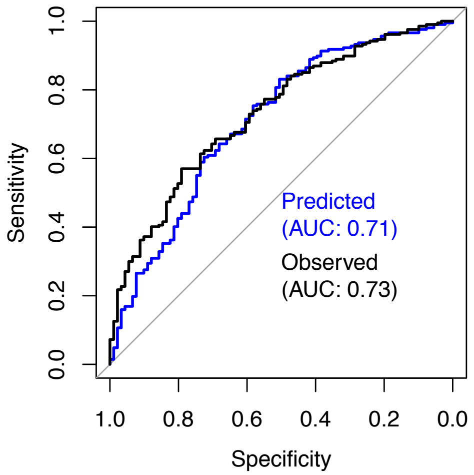


**Figure S5. Prediction accuracy of 5-year survival (Yes/No).** We compared prediction accuracy of 5-year survival using predicted gene expression vs observed gene expression.

**Table S1.** Confusion Matrix: Predicting breast cancer sub-type using (a) gene expression predicted by geneEXPLORER (b) Observed gene expression

| | | Predicted | | | | | | Predicted | | | | |
|---|---|---|---|---|---|---|---|---|---|---|---|---|
| | | Basal | Her2 | LumA | LumB | Normal | | Basal | Her2 | LumA | LumB | Normal |
| True | Basal | 84 | 1 | 1 | 0 | 1 | Basal | 85 | 1 | 0 | 1 | 0 |
| | Her2 | 3 | 22 | 0 | 6 | 0 | Her2 | 0 | 22 | 1 | 8 | 0 |
| | LumA | 0 | 0 | 252 | 27 | 9 | LumA | 0 | 1 | 260 | 17 | 10 |
| | LumB | 0 | 0 | 49 | 78 | 0 | LumB | 0 | 1 | 33 | 93 | 0 |
| | Normal | 2 | 1 | 8 | 0 | 76 | Normal | 2 | 1 | 7 | 0 | 77 |

A. Predicted gene expression
Misclassification error: 0.174

B. Observed gene expression
Mis-classification error: 0.134
